# Genetic analysis and ecological niche modeling delimit species boundary of the Przewalski’s scorpion (Scorpiones: Buthidae) in arid Asian inland

**DOI:** 10.1101/652024

**Authors:** Xue-Shu Zhang, Gao-Ming Liu, Yu Feng, De-Xing Zhang, Cheng-Min Shi

**Affiliations:** State Key Laboratory of Integrated Management of Pest Insects and Rodents, Institute of Zoology, Chinese Academy of Sciences, Beijing 100101, China; CAS Key Laboratory of Genomic and Precision Medicine, Beijing Institute of Genomics, Chinese Academy of Sciences, Beijing 100101, China

**Keywords:** *Mesobuthus*, species complex, mitochondrial DNA, ecological niche modeling, distribution range

## Abstract

Reliable delimitation of venomous scorpions is not only consequential to toxicological studies but also instructive to conservation and exploration of these important medical resources. In the present study, we delimited species boundary for the the Przewalski’s scorpion from arid northwest China through a combined approach employing phylogenetic analysis, ecological niche modeling and morphological comparison. Our results indicate that the the Przewalski’s scorpion represent an independent taxonomic unit and should be recognized as full species rank, *Mesobuthus przewalsii* **stat. n.**. This species and the Chinese scorpion *M. martensii* represent the eastern members of the *M. caucasicus* species group which manifests a trans-Central Asia distribution across the Tianshan Mountains range. We also discussed the likely geographic barrier and climatic boundary that demarcate distributional range of the the Przewalski’s scorpion.

## 1 Introduction

Reliable delimitation of species is crucial as species are fundamental units of ecology, biodiversity, macroevolution and conservation biology (Sites & Marshall 2004). While different ideas are prominent in various naturalists’ minds when speaking of species, a common element is that species are ‘separately evolving metapopulation lineages’ (De Queiroz 2007). Following such a unified species concept, species can be delimited in an integrative way based on operational criteria that reflect contingent, but not necessary, properties of the common elements, such as genetic divergence and ecological distinctiveness besides morphological difference that are traditionally used to recognize species (Leaché et al. 2009; Padial et al. 2010).

Genetic and ecological properties appear to be relevant to species delimitation for groups that suffer taxonomic confusion in the traditional morphological framework. Genetic approaches, especially those based on DNA sequences which provide highly variable and stable characters, have revolutionized our ability for species delimitation (Tautz et al. 2003; Godfray et al. 2004; Savolainen et al. 2005; Vogler & Monaghan 2007) and become an indispensable means for resolving various kinds of taxonomic problems. Ecological approaches, particularly the ecological niche modeling (ENM), associate environmental variables with known species’ occurrence localities to define abiotic conditions within which populations can survive (Guisan & Thuiller 2005). Thus ENM can provide compelling evidence for geographic isolation between allopatric populations, which has important practical implications for species delimitation (Wiens & Graham 2005). It also helps delineate spatial distribution boundaries based on the abiotic factors even when species occurrence is known from very limited points (Pearson et al. 2007). Clearly, combining analysis of DNA sequences and ENM together with the traditional morphological evidences would warrant robust delimitation of species boundaries. However, such an integrated approach remains to be fully employed in the taxonomic studies of arachnids, particularly of scorpions which are known for their morphological conservatism and taxonomical difficulty.

Scorpions of the *Mesobuthus* Vachon 1950 (Scorpiones: Buthidae) are venomous to human beings and have subjected to intensive toxicological studies (Goudet et al. 2002; Cao et al. 2006; Zhu et al. 2012; Diego-García et al. 2013). Despite their medical importance and pharmaceutical significance, however, taxonomic status of species in this genus remains to be critically and robustly delimitated. Scorpions of this genus occur widely in the temperate Palaearctic region from Balkan in the west to coastal China in the east (Shi & Zhang 2005; Shi et al. 2007, 2013). A plethora intraspecific units, such as subspecies, forms and types, have been recognized within wide-ranging species (Fet et al. 2000). Recent treatments, particularly analyses of DNA sequences, of widespread polytypic species have led to recognition more than ten new species (Gantenbein et al. 2000; Fet et al. 2018; Mirshamsi et al. 2010, 2011). It has been also demonstrated that climatic niche is an important determinant of geographic distribution of *Mesobuthus* scorpions (Shi et al. 2007, 2015). Species differentiation appears to associate with significant divergence in their ecological niches (Mirshamsi 2013). Thus, we expect that an integrated approach combining DNA sequence analysis and ecological niche modeling will bring to a robust delimitation of species in this group.

The aim of the present study is thus to clarify the taxonomic status of *Mesobuthus* species from China. Up to date, six *Mesobuthus* species have been reported from China, viz. *M. bolensis, M. caucasicus, M. eupeus, M. karshius, M. longichelus* and *M. martensii*. Three of them (*M. caucasicus, M. eupeus*, and *M. martensii*) are geographically widespread, each having two subspecies been recorded in China. Other three species (*M. bolensis, M. karshius* and *M. longichelus*) were described recently and narrowly distributed (Sun & Sun 2011; Sun & Zhu 2010; Sun et al. 2010). The species *M. caucasicus* is one of the most morphologically diverse and geographically wide-spread species in the genus. It ranges from Caucasian Mountains in the west to the northwest China in the east. Historically, six subspecies have been recognized (Fet et al. 2000). Recently revision based on morphology and mitochondrial DNA sequences revealed that this taxon represents a species complex; four subspecies have been elevated to species rank and six additional new species were described from Central Asia (Fet et al. 2018). However, the only subspecies which occurs to the east of the Tianshan Mountains and the Pamir Plateau (Birula 1897), the Przewalski’s scorpion *M. caucsicus przewalsii* was not examined. Here we focus specially on the Przewalski’s scorpion, of which taxonomic status is unclear and has never been subjected to genetic and ecological assessment. We present our results from morphological examination, genetic analysis and ecological niche modeling that clarify the taxonomic status of the Przewalski’s scorpion and its geographical range.

## 2 Materials and Methods

### 2.1 Sampling and morphology

Samples were collected during filed surveys of *Mesobuthus* scorpion diversity and geographic distribution by a stone-rolling method in daytime or UV-light searching at night. Special efforts were taken to survey the region around the type localities in the southeast part of the Tarim Basin. The original descriptions of two type localities (i.e. near Lob-nor, Ruoqiang and Oasis Cherchen, Qiemo) are very vague (Birula, 1897). We surveyed along the Cherchen river and successfully collected specimens from Qiemo and Ruoqiang. Although we did not reach the Lob-nor, our sampling sites already surrounded the region. The sampling localities were geo-positioned using a GPS receiver (Garmin International). Morphological observation was performed under a Nikon SMZ1500 stereomicroscope. Samples are preserved in 99.7% ethanol and deposited at Institute of Zoology, Chinese Academy of Sciences, Beijing.

### 2.2 Material examined

CHINA: *Xinjiang:* 3 ♂, 4 ♀, Tuokexun, 42°47’N, 88°40’E, 12 September 2015, C. -M. Shi; 1 ♂, 8 ♀, Turpan, 42°57’N, 89°06’E, 20 September 2017, C.-M. Shi & G.-M. Liu; 2 ♂, 3 ♀, Hami, 42°52’N, 93°26’E, 14 September 2015, X .-G. Guo; 3 ♀, Hami, 42°41’N, 93°27’E, 22 September 2017, C.-M. Shi & G.-M. Liu; 3 ♂, 1 ♀, Kuche, 41°44’N, 82°55’E, 20 August 2018, C.-M. Shi; 3 ♀, Tiemenguan, 41°47’N, 86°11’E, 21 September 2017, G.-M. Liu & Y. Feng; 1 ♂, 3 ♀, Shanshan, 42°51’N, 90°13’E, 11 September 2015, C. -M. Shi; 9 ♂, 8 ♀, Tumushuke, 39°53’N, 78°55’E, 24 August 2018, C.-M. Shi; 1♀, Tarim, 40°38’N, 84°18’E, 21 June 2015, X.-G. Guo; 5 ♂, 3 ♀, Qiemo, 38°15’N, 85°32’E, 26 August 2019, C. -M. Shi & Y. Feng; 2 ♂, 7 ♀, Ruoqiang, 39°01’N, 88°10’E, 27 August 2019, C.-M. Shi & Y. Feng; *Gansu*: 1 ♂, Dunhuang, 40°18’N, 94°44’E, 23 September 2017, C.-M. & G.-M. Liu; 2 ♂, 3 ♀, Guazhou, 40°29’N, 96°02’E, 16 September 2015, C.-M. Shi; *Inner Mongolia*: 1 ♂, 1 ♀, Ejina, 42°04’N, 101°12’E, 5 September 2015, C.-M. Shi.

### 2.3 Molecular phylogenetic analysis

One specimen from each of collection site was used for sequencing the mitochondrial cytochrome c oxidase subunit I (mtCOI). Genomic DNA was extracted from preserved tissues using a modified phenol-chloroform extraction procedure (Zhang & Hewitt 1998). The primers, PCR profiles and sequencing protocols followed Shi et al. (2013). The unique haplotypes have been deposited in GenBank under accession numbers xxxxxxxx-xxxxxxxx. Sequences for other species of the *M. caucasicus* complex recognized by Fet et al. (2018) were downloaded from GenBank. We also included six sequences which represented the major mitochondrial lineages of the Chinese scorpion *M. martensii* (Shi et al. 2013), and seven sequences of the mottled scorpion *M. eupeus monglicus* from the north side of the Tianshan Mountains (Table 1). Sequences were aligned together with the sequences generated in the present study using CLUSTAL X 1.83 (Thompson et al. 1997) and further inspected by eye. Phylogenetic analyses were performed using both maximum likelihood (ML) and Bayesian methods. ML analysis was carried out using IQ-TREE v1.6.10 (Nguyen et al. 2015) with DNA substitution model selected by ModelFinder (Kalyaanamoorthy et al. 2017). Branch supports were obtained with the ultrafast bootstrap with 1000 replicates (Hoang et al. 2018). Bayesian analysis was carried out with MrBayes 3.2 (Ronquist et al. 2012). Analyses were initiated with random starting trees and run for 2×10^6^ generations with four Markov chains employed. Trees were sampled every 200 generations and the ‘temperature’ parameter was set to 0.2. The first 25% trees were discarded as burn-in after checking for stationary and the convergence of the chains. The approximately unbiased (AU) test (Shimodaira 2002) was used to evaluate the alternative phylogenetic hypotheses. Genetic distances between morphologically and/or phylogenetically recognized species were calculated using Kimura 2-parameter model (K2P distance) with MEGA 5.05 (Tamura et al. 2011).

**Table 1.**
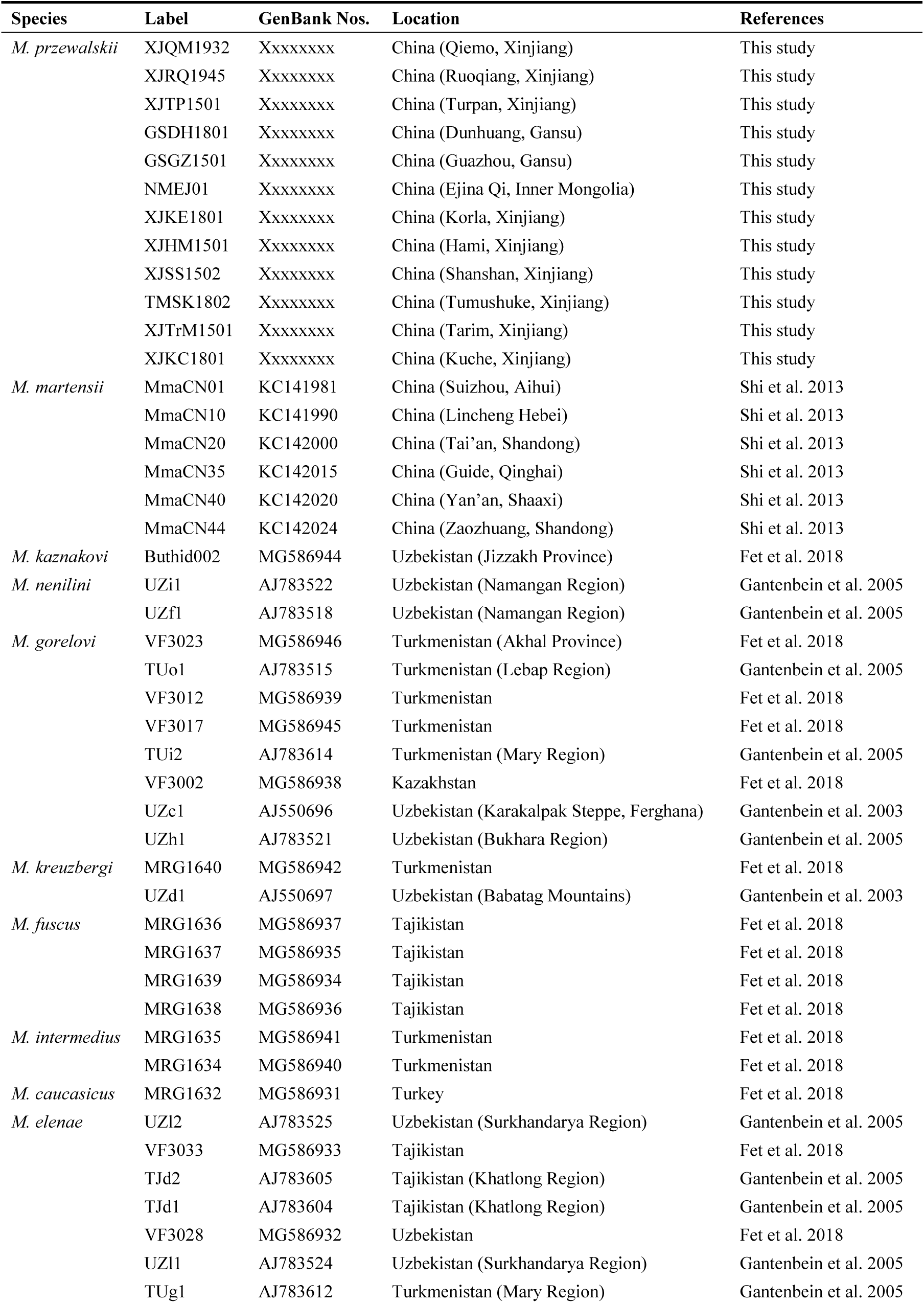
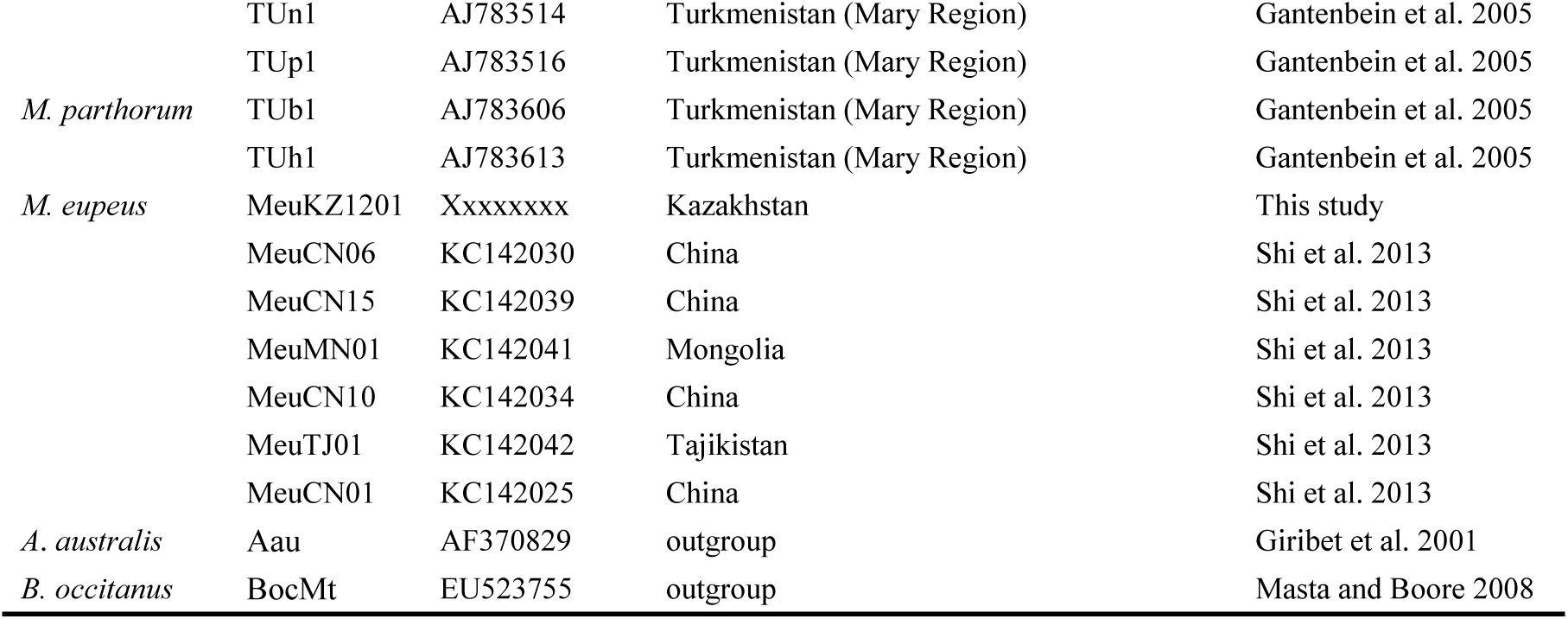
Genbank accession numbers of *Mesobuthus* scorpions and outgroup sequences.

### 2.4 Ecological niche modeling

We predicted potentially suitable distribution area through ecological niche modeling (ENM). ENM was performed using MaxEnt version 3.4.1 (Phillips & Dudik 2008) based on scorpion occurrence points and bioclimatic variables. For the Przewalski’s scorpion, a total of 12 GPS points were recorded during field survey for this study and additional 19 occurrence records were georeferenced from literatures. The bioclimatic variables were download from the WorldClilm database (http://www.worldclim.org/). These climatic variables represent a set of measurements that summarize temperature and precipitation at a 2.5 arc-minute resolution (c. 5×5 km). We masked the climatic variable to known range of the *M. caucasicus* complex, spanning from 30 to 50 °N and from 40 to 105 °E, to avoid sampling unrealistic background data and thus inflating the strength of predictions. We removed highly correlated (Pearson’s correlation, |r| ≥ 0.80) climatic variables before performing ENM. MaxEnt was run with a convergence threshold of 10^−5^ and maximum number of iterations of 10,000 with cross validation. Model performance was assessed via the area under the ROC (receiver operating characteristic) curve (AUC) statistic and the importance of variables was assessed by jackknife tests. We employed the maximum training sensitivity plus specificity threshold for converting continuous models to binary predictions. We did not confine the background sampling of ENM within the minimum convex polygon of the occurrence points so that the suitable distribution area was slightly over-predicted. Ecological niche models for the geographically neighboring species, the Chinese scorpion *M. martensii* (Shi et al. 2007) were projected together with the Przewalski’s scorpion to check range overlap visually. To test whether ecological niche of the Przewalski’s scorpion overlaps with other species of the *M. caucasicus* complex, we also performed independent ENM for these species collectively since the phylogenetically validated occurrence data for each individual species was very few. A total 74 occurrence points for nine species (Fet et al. 2018), all of which occur to the west of the Tianshan Mountains, were used to construct a single distribution model. Such a lumping prediction will cause over-prediction for individual species due to sampling background points in a wider and unrealistic region for the relevant species. Given the over-prediction of both models, we considered they gave over-estimate of the degree of distributional overlap, in another words, a conservative prediction of distribution separation.

## 3 RESULTS

### 3.1 Taxonomy

**Family** Buthidae C. L. Koch, 1837

**Genus** *Mesobthus* Vachon, 1950

***Mesobuthus przewalskii*** (Birula, 1897) **stat. n.**

*Buthus caucasicus przewalskii*, Birula, 1897:337-338.

*Buthus przewalskii*: Kishida, 1939:44.

*Mesobuthus caucasicus intermedius*: Vachon, 1958: 150, Fig. 31.

*Olivierus caucasicus intermedius*: Farzanpay, 1987: 156; Fet et al. 2000: 191; Zhu et al. 2004:113.

*Mesobuthus caucasicus przewalskii*: Shi & Zhang 2005:475; Sun & Zhu 2010:4-5; Sun & Sun 2011:60-61; Di et al. 2015:111

**Type locality.**—CHINA (near Lob-nor, Ruoqiang and Oasis Cherchen, Qiemo in the east edge of the Tarim Basin, Xinjiang)

**Distribution.**—CHINA (Tarim Basin, Xinjiang; Ejina Qi, Inner Mongolia; Guazhou and Dunhuang, Gansu)

**Diagnosis.**—Total length 50-61 mm in adult males, 53-75 mm in females (Figs. 1-4). Pedipalp femur with 4-5 granulate carinae, patella with 8 granulate or smooth carinae, manus with irregular netlike dark pigments; chela slender with Cl/Cw = 3.98±0.38 (Cl = chela length, Cw = chela width, mean ± SD, n = 43) for the female and 3.07 ± 0.28 (n=28) for the male. Trichobothrium *db* on fixed finger of pedipalp situated between trichobothria *esb* and *est*, near to *est*. For both sexes, movable and fixed fingers of pedipalps with 11 and 10 cutting rows of denticles, respectively (Figs. 5-6); movable fingers with 5 terminal denticles. Pectinal teeth number 15-19 in females and 19-23 in males (Figs. 7-8). Seventh sternite bears 4 marked granulate carinae. Metasomal segments I-V sparsely hirsute with 10-8-8-8-5 complete carinae; ventrolateral carinae strong, serrate, becoming stronger gradually from anterior to posterior. Lateral and dorsal surfaces of metasomal segment V with sparse granules and ventral with sparse large granules. The telson sparsely hirsute, bumpy; aculeus about equal to a half of telson length. Leg femur and patella with carinae; long tibial spurs present on legs III and IV; pedal spur with solitary setae; tarsomeres with two rows of setae on ventral surface and numerous macrosetae on the other surfaces. Telotarsus III ventrally bear two rows of setae in which every contains not more than 13 setae in both sexes.

**Figures 1-8.**
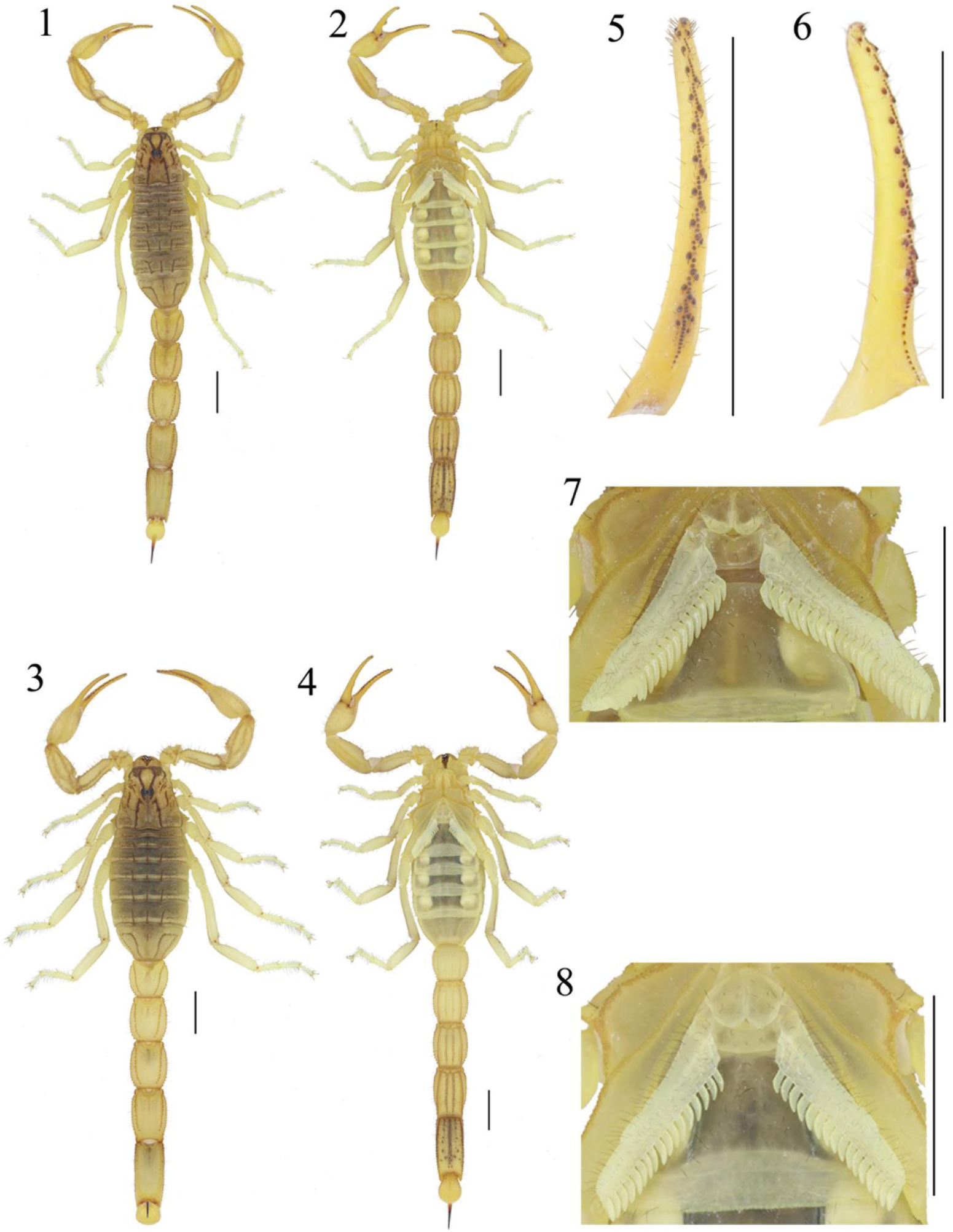
**1-2, 5-7. *Mesobuthus przewalsii* stat. n., male from Qiemo county, Xinjiang**: 1. Dorsal view. 2. Ventral view. 5. Dentition of pedipalp chela movable finger. 6. Dentition of pedipalp chela fixed finger. 7. Ventral aspect of genital operculum and pectines. **3-4, 8. *Mesobuthus przewalsii* stat. n., female from Qiemo county, Xinjiang**: 3. Dorsal view. 4. Ventral view. 8. Ventral aspect of genital operculum and pectines. Scale bars = 5.0 mm.

### 3.2 Genetic divergence and phylogenetic relationship

A best-fit model of TIM3+F+R3 was selected by ModelFinder. Based on this evolution model, IQ-TREE inferred a ML tree with optimal log-likelihood of −3993.82 from mtCOI DNA sequences (Fig. 9). In the ML tree, most species of the *M. caucasicus* complex formed strongly supported (bootstrap support values ≥ 95) monophyletic clade except for *M. elenae* which appeared paraphyletic with respect to *M. parthorum* (Fig. 9). Robustness of the ML tree was tested against a constrained tree in which *M. elenae* and *M. parthorum* were forced to be reciprocal monophyly with the AU test. The ML tree has a high value of log likelihood than the constrained tree (−3998.73) and was preferred by the AU test but not significantly (*P*_-AU_ = 0.67). The relationship between the two species remains to be resolved in future. Bayesian MCMC sampling was converged after 2×10^6^ generations as indicated by the value of the potential scale reduction factor (PSRF) approaching 1.00 and the average standard deviation of split frequencies less than 0.0075. All clades corresponding to species recognized in the ML tree were strongly supported (posterior probability = 1.00) in the Bayesian majority role consensus tree (Fig. S1). However, the inter relationships among species of the *M. caucasicus* complex were unresolved, forming a multifurcating clade harboring *M. martensii* and all member of the *M. caucasicus* complex except *M. elenae* and *M. parthorum* which formed a paraphyletic clade as in the ML tree. All samples for the Przewalski’s scorpion clustered in a fully supported monophyletic clade (100/1.00), which was clearly separated from other members of the complex. The Przewalski’s scorpion appeared to be most closely related to a lineage composed of *M. nenilini and M. kaznakovi* in the ML tree but such relationship was not supported by the Bayesian tree. Interestingly, the Chinese scorpion *M. martensii* was nested within the clade for the *M. caucasicus* complex (Fig. S1).

**Figure 9.**
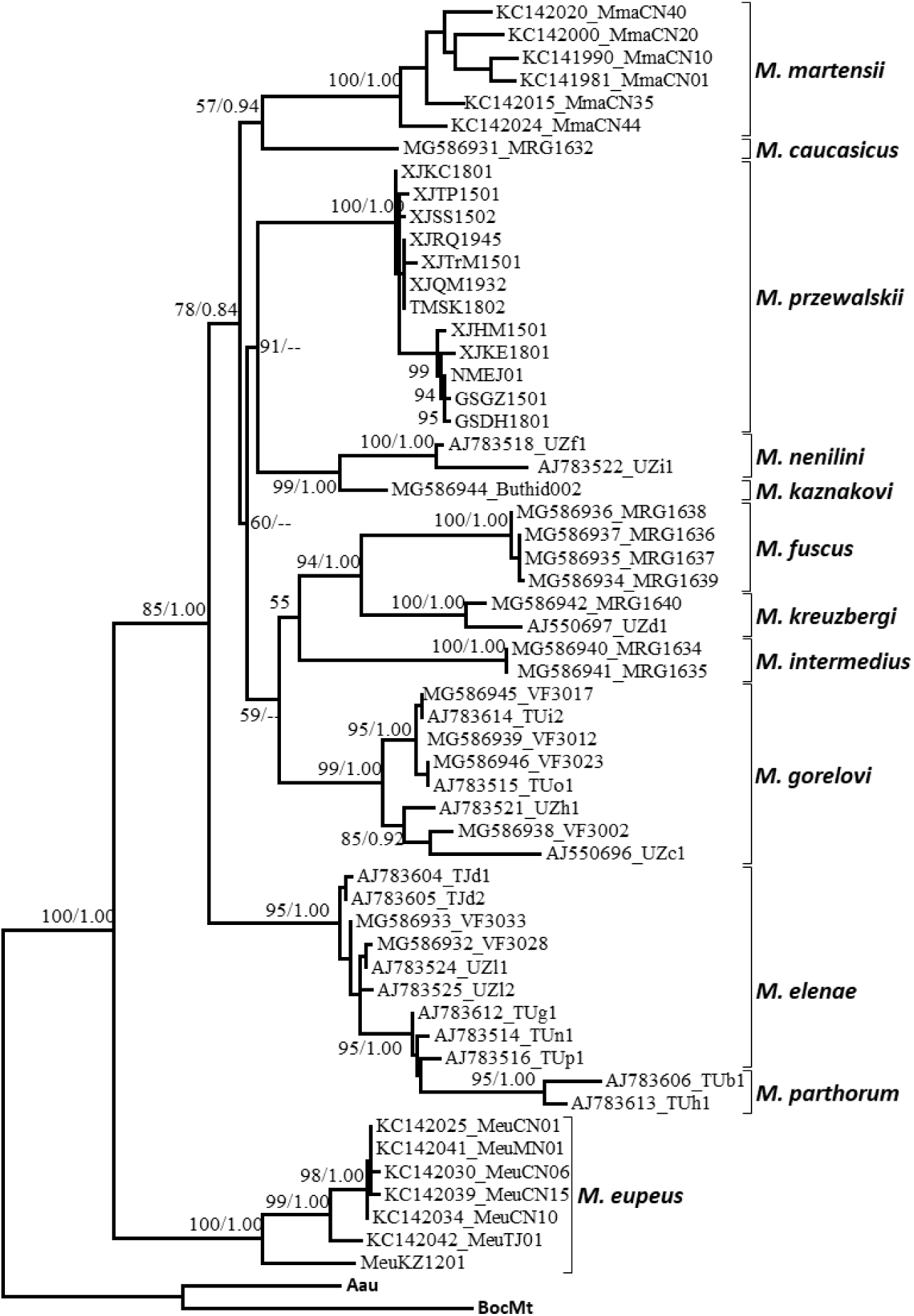
Phylogeny the *Mesobuthus caucasicus* complex reconstructed using mitochondrial DNA sequences. The Przewalski’s scorpion (*M. przewalskii*) is deeply diverged from other species and the Chinese scorpion (*M. martensii*) belongs to the species complex. Node supports are shown by the exact numbers for bootstrapping probabilities from 1000 replicates and the posterior probability.

The interrelationships among species of the *M. caucasicus* complex is shown by phylogenetic network to illustrate the phylogenetic uncertainty (Fig. 10). The result was completely in congruence with the maximum likelihood tree, for which poorly supported internodes were indicated by reticulations. Although the interrelationships between species is poorly resolved, no reticulations have occurred in the most recent common ancestors for each species of the complex. The Przewalski’s scorpion is clearly diverged from other member of the species complex and warrants a species rank. The divergence of the Chinese scorpion *M. martensii* is comparable to the divergences among the members of the species complex. The Estimates of net evolutionary divergences in the mtCOI sequences among species are shown in Table 2. The interspecies genetic distances (based on K2P model) among member of the *M. caucasicus* complex ranges from 3.96% (*M. elenae* vs. *M. parthorum*) to 10.29% (*M. intermedius* vs. *M. fuscus*). The genetic distances between the Przewalski’s scorpion other species of the complex ranges 7.30% to 9.41%. It appeared that the Przewalski’s scorpion was diverged more from other member of the *M. caucasicus* species complex (≥7.30%) than from the Chinese scorpion *M. martensii* (6.10%).

**Table 2.** Estimates of net evolutionary divergence in the mtCOI sequences between species of *Mesobuthus caucasicus* complex. Show here are the mean ± standard error of net genetic distances calculated using the Kimura 2-parameter model (lower triangle). Entries on the diagonal present means of within species genetic distances and their standard errors.

|  | *M. przewalskii* | *M. martensii* | *M. gorelovi* | *M. kaznakovi* | *M. elenae* | *M. kreuzbergi* | *M. caucasicus* | *M. nenilini* | *M. parthorum* | *M. intermedius* | *M. fuscus* |
| --- | --- | --- | --- | --- | --- | --- | --- | --- | --- | --- | --- |
| *M. przewalskii* | **0.0118 ± 0.0035** |  |  |  |  |  |  |  |  |  |  |
| *M. martensii* | 0.0610 ± 0.0107 | **0.0348 ± 0.0060** | | | | | | | | | |
| *M. gorelovi* | 0.0730 ± 0.0146 | 0.0682 ± 0.0132 | **0.0264 ± 0.0052** | | | | | | | | |
| *M. kaznakovi* | 0.0752 ± 0.0138 | 0.0762 ± 0.0131 | 0.0733 ± 0.0157 | **N.A** | | | | | | | |
| *M. elenae* | 0.0772 ± 0.0118 | 0.0593 ± 0.0099 | 0.0669 ± 0.0110 | 0.0662 ± 0.0123 | **0.0186 ± 0.0040** | | | | | | |
| *M. kreuzbergi* | 0.0826 ± 0.0138 | 0.0701 ± 0.0119 | 0.0764 ± 0.0155 | 0.0822 ± 0.0149 | 0.0794 ± 0.0144 | **0.0272 ± 0.0077** | | | | | |
| *M. caucasicus* | 0.0819 ± 0.0154 | 0.0715 ± 0.0133 | 0.0751 ± 0.0148 | 0.0784 ± 0.0150 | 0.0856 ± 0.0113 | 0.0884 ± 0.0166 | **N.A** | | | | |
| *M. nenilini* | 0.0864 ± 0.0153 | 0.0749 ± 0.0142 | 0.0709 ± 0.0140 | 0.0411 ± 0.0096 | 0.0619 ± 0.0120 | 0.0855 ± 0.0150 | 0.0920 ± 0.0156 | **0.0319 ± 0.0090** | | | |
| *M. parthorum* | 0.0898 ± 0.0149 | 0.0706 ± 0.0124 | 0.0547 ± 0.0110 | 0.0735 ± 0.0140 | 0.0396 ± 0.0103 | 0.0768 ± 0.0155 | 0.0844 ± 0.0146 | 0.0795 ± 0.0161 | **0.0727 ± 0.0129** | | |
| *M. intermedius* | 0.0941 ± 0.0158 | 0.0728 ± 0.0128 | 0.0666 ± 0.0121 | 0.0995 ± 0.0179 | 0.0923 ± 0.0134 | 0.0888 ± 0.0151 | 0.0963 ± 0.0154 | 0.0992 ± 0.0167 | 0.0801 ± 0.0131 | **0.0000 ± 0.0000** | |
| *M. fuscus* | 0.0803 ± 0.0147 | 0.0720 ± 0.0121 | 0.0735 ± 0.0149 | 0.0758 ± 0.0150 | 0.0854 ± 0.0143 | 0.0715 ± 0.0142 | 0.0857 ± 0.0154 | 0.0831 ± 0.0139 | 0.0865 ± 0.0163 | 0.1029 ± 0.0169 | **0.0011 ± 0.0011** |
| *M. eupeus* | 0.1203 ± 0.0180 | 0.1068 ± 0.0162 | 0.0906 ± 0.0142 | 0.0978 ± 0.0142 | 0.1039 ± 0.0154 | 0.1039 ± 0.0129 | 0.1167 ± 0.0167 | 0.0999 ± 0.0156 | 0.0801 ± 0.0129 | 0.1257 ± 0.0192 | 0.1101 ± 0.0159 |

**Figure 10.**
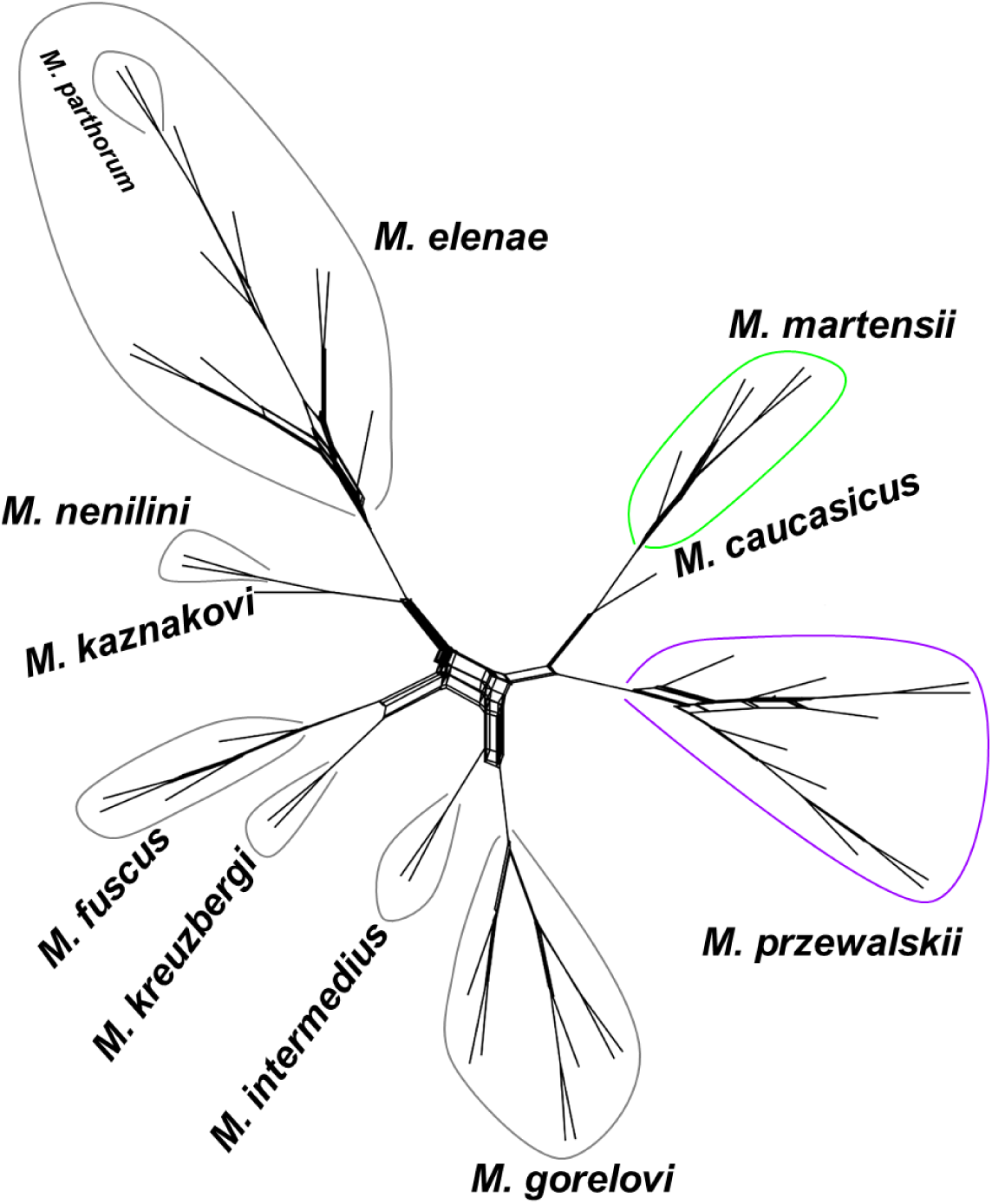
Phylogenetic network for the *Mesobuthus caucasicus* species complex. Although the interrelationships between species is poorly resolved, no reticulations have occurred in the most recent common ancestors for each species. The Przewalski’s scorpion is clearly diverged from other member of the species complex and warrants a species rank. The divergence of the Chinese scorpion *M. martensii* is comparable to the divergences among the members of the species complex.

### 3.3 Geographic distribution and ecological niche modeling

We assembled a total of 31 geographic points, 12 recorded from field survey and 19 georeferenced from literatures, for occurrence of the Przewalsk’s scorpion. These occurrence points cover a large geographic space, stretching about 2000 km from east to west. All occurrence sites are to the south of the Tianshan Mountains range and mostly in the Tarim Basin. After removing highly correlated (|r| ≥ 0.80) climatic variables, six variables were used in ecological niche modeling. These include Bio3 = isothermality, Bio8 = Mean Temperature of Wettest Quarter, Bio9 = Mean Temperature of Driest Quarter, Bio13 = Precipitation of Wettest Month, Bio14 = Precipitation of Driest Month, and Bio15 = Precipitation Seasonality. We also performed ENM using only the 12 occurrence sites from which samples were sequenced for phylogenetic analysis. The predicted potential distribution areas almost overlapped that predicted with all occurrence sites. Thus, we only report the ENM results based on the full distributional records here. The relative contributions of these six climatic variables to the model ranged from 1.3% (Bio3) to 54.0% (Bio13). The ENM performed very well with an AUC value of 0.976.

The predicted suitable distribution areas for the Przewalski’s scorpion are shown in Figure 11. The entire Tarim basin and adjacent Gobi regions are suitable for survival of *M. przewalsii*, however, no areas to the west of the Tianshan Mountains and the Pamir Plateau is predicted suitable. On the contrary, the predicted suitable distribution areas for all other nine species of the *M. caucasicus* complex lumped together effectively restricted to the west of the Tianshan Mountains and the Pamir Plateau (Fig. 11). The predicted distribution ranges for these scorpions were clearly separated by unsuitable areas composed of the Tianshan Mountains from the suitable range of the Przwewalsi’s scorpion in the Tarim Basin. No overlap was observed in potential distribution range between the Przwewalsi’s scorpion and other species of the complex (Fig. 11). It appears that the unsuitable Tianshan Mountains and the Pamir Plateau form unsurmountable barriers that have isolated the Przwewalsi’s scorpion in Tarim from other species in Central Asia. The large suitable areas were predicted in the Junggar basin to the north of the Tianshan Mountains. However, our field survey suggested that this species does not occur in these regions. On the contrary, another species *M. eupeus mongolicus* is very common and these regions represent the core distribution range of the later (Shi et al. 2015). These areas may represent artificial suitable niches over-predicted by the abiotic model or represent real suitable niches that could not have been explored by the Przwewalsi’s scorpion due to the existence of geographic barriers, i.e. the Tianshan Mountains. There are limited overlaps in predicted suitable distribution areas between *M. przewalsii* and *M. martensii* along the northeast edge of the Qinghai-Tibet Plateau. Field survey also implicates that contact between the Przwewalsi’s scorpion and the *M. martensii* is unlikely. There is at least 500 km distribution gap between these two species along the Hexi corridor, which is currently distributed by *M. eupeus mongolicus* (Shi et al. 2015). Taken together, we define the Tainshan Mountains as the northern boundary, the western entrance of the Hexi corridor as the eastern boundary, the Pamir as the western boundary and the Qing-Tibet Plateau as the southern boundary, respectively, for the Przwewalsi’s scorpion.

**Figure 11.**
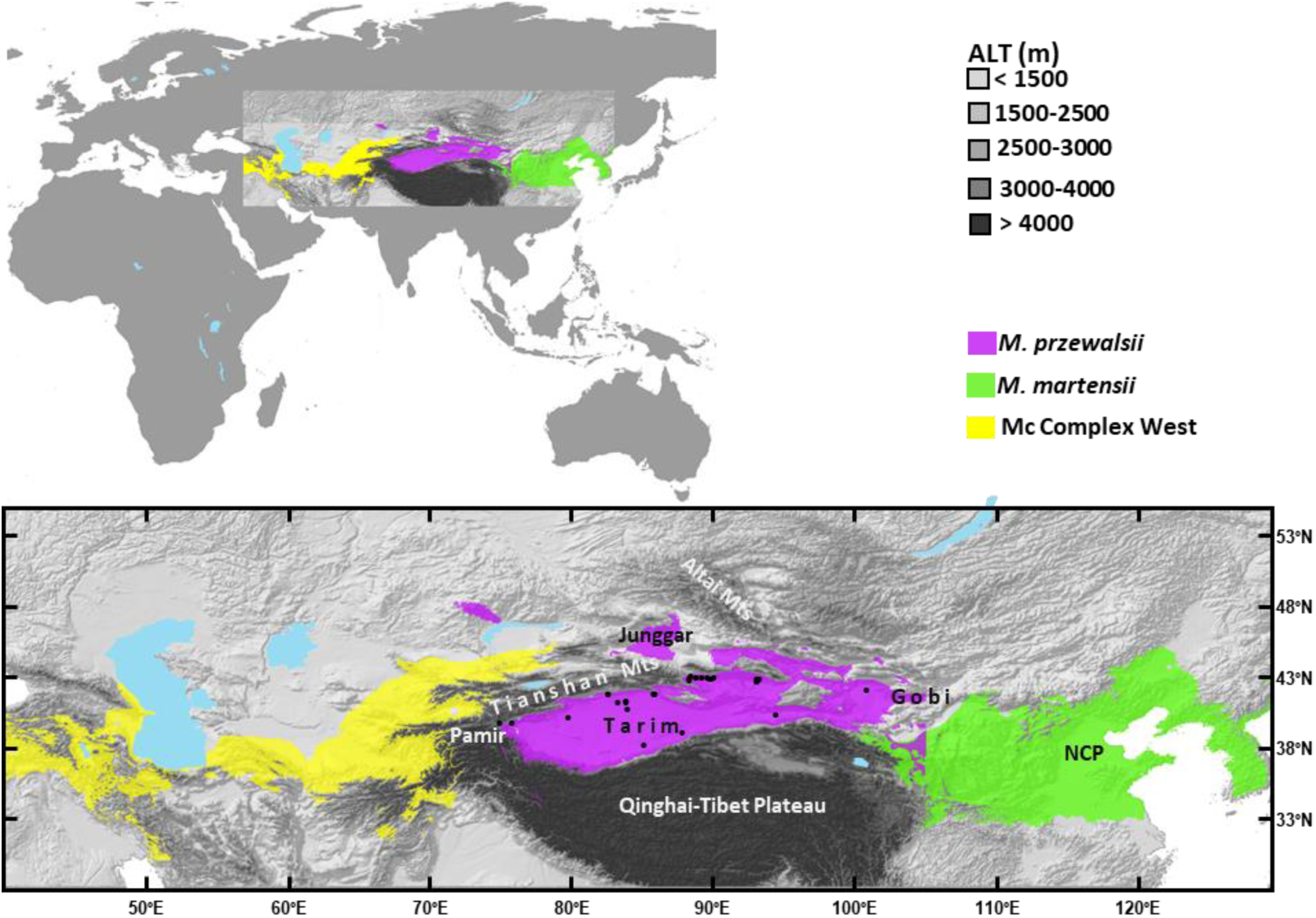
Ecological niche models of *Mesobuthus* scorpions. Potential distribution areas for the Przewalski’s scorpion (*M. przewalsii*, purple) is shown together with the Chinese scorpion (*M. martensii*, light green) and other species of the *M. caucasicus* complex (Mc Complex West, yellow). The entire Tarim Basin and adjacent Gobi region are suitable for survival of *M. przewalsii*. No area to the west of the Tianshan Mts and the Pamir Plateau is suitable for *M. przewalsii*, and similarly no area to the east of the Tianshan Mts and Pamir Plateau is suitable for other species of the *M. caucasicus* complex. There are overlaps in predicted suitable distribution areas between *M. przewalsii* and *M. martensii* along the northeast edge of the Qinghai-Tibet Plateau. The suitable areas in the Junggar Basin and two the north of the Tianshan Mts are due to over prediction of the model, because *M. przewalsii* does not occur in these regions. Ecological niche model for *M. martensii* was adopted from Shi et al. 2007.

## 4 Discussion

### 4.1 Multifaceted recognition of the Przewalski’s scorpion

According the unified species concept, species represent separately evolving metapopulation lineages (De Queiroz 2007), which possess many contingent, but not necessary, properties making them be reciprocally monophyletic, ecologically divergent, or morphologically distinctive (Leaché et al. 2009). In the present study, we adopt the unified species concept and use these contingent properties as the operational criteria to delimit species for the Przewalski’s scorpion, *M. przewalskii*.

Morphologically, the Przewalski’s scorpion can be distinguished from other related species by several stable diagnostic characters. The Przewalski’s scorpion is most similar to *M. intermedius* (Birula 1897; Sun & Zhu 2010). However, two species can be distinguished by the number of pectinal teeth. Pectines of the Przewalski’s scorpion have 15-19 teeth in females and 19-23 teeth in males, while those of *M. intermedius* have 20-25 teeth in females and 26-30 teeth in males, respectively (Sun & Zhu 2010). In comparison with two parapatric species, the movable and fixed fingers have 11 and 10 oblique rows of granules respectively, while both the movable and fixed fingers of *M. martensii* have 12-13 rows of oblique granules, and those of *M. eupeus mongolicus* have 11 and 10 rows, respectively (Shi et al. 2007). The Cl/Cw ratio of pedipalp chela of the Przewalski’s scorpion is 3.98±0.38 for the female and 3.07 ± 0.28 for the male, while *M. martensii* has Cl/Cw ratio of 4.45±0.23 for the female and 3.71 ± 0.24 for the male (Shi et al. 2007).

Genetically, the mitochondrial DNA data revealed that the Przewalski’s scorpion represents monophyletic taxonomic unit. In the phylogenetic analyses of mtCOI DNA sequences, all samples of the Przewalski’s scorpion clustered together and formed a strongly supported monophyletic clade in both ML and Bayesian phylogenies (Figs. 9 & S1). In addition, we found deep genetic divergences of 6.1-9.4% between the Przewalski’s scorpion and other members of the *M. caucasicus* complex. Assuming a divergence rate of 1.7% for *Mesobuthus* scorpions (Shi et al. 2013), this genetic distance corresponds to a divergence time of 3.6-5.5 million years. Such a long-term period would allow enough time for sorting of ancestral polymorphisms so that species become reciprocally monophyletic.

Ecologically, our results of ENM suggested that the Przewalski’s scorpion occupies a distinctive geographic range. The core potential distribution areas for the Przewalski’s scorpion are confined in the Tarim Basin and the surrounding high mountain ranges are not suitable for survival of scorpions (Fig. 11). This observation implies that the high mountains, particularly the Tianshan, have constituted an important geophysical barrier. The central range of Tianshan Mountains appeared impermeable and isolated the Przewalski’s scorpion from other Central Asian species of the *M. caucasicus* complex in the west and the mottled scorpion *M. eupeus mongolicus* in the north. On the contrary, there appears no significant physical barrier that would isolate the Przewalski’s scorpion from *M. martensii* and *M. eupeus mongolicus* outside of the Tarim Basin in the east. Consistent with studies on other congeneric species (Mirshamsi 2013; Shi et al. 2007, 2015), it appeared that climatic variables were also a determinative factor in defining geographic distribution of the Przewalski’s scorpion. This point was particularly supported by its eastern distribution boundary, where no prominent geophysical barrier existed. Its potential distribution range was clearly separated from that of its geographically neighboring species, *M. eupeus mongolicus* in the north (Shi et al. 2015) and *M. martensii* in the east (Shi et al. 2007). Historically, the Prezwalski’s scorpion was first described from the eastern edge of the Tarim Basin, near Lob-nor and oasis of the Cherchen river (Birula, 1897). Earlier reports suggested that this species might wide spread in Xinjiang, northwest China (Sun & Zhu 2010; Sun & Sun 2011) and may also occur in Mongolia, Tajikistan and Uzbekistan (Fet et al. 2000). However, recent revisions (Fet et al. 2018; Kovařík 2019) and our result of ecological niche modeling implied that its occurrence to the north (i.e. the Junggar Basin) and west (i.e. Tajikistan and Uzbekistan) of the Tianshan Mountains range can be tentatively excluded (Fig. 10). Recent field surveys also failed to conform the presence of *M. przewalskii* in Mongolia (Shi et al. 2015) and it appeared that earlier Mongolian records were originated from present-day China (Heddergott et al. 2016). Currently confirmed distribution of *M. przewalskii* was limited to the arid northwest China.

In summary, all above evidences indicate that the Przewalski’s scorpion is morphologically distinctive, genetically differentiated, phylogenetically independent and ecologically divergent from other *Mesobuthus* species. These contingent properties collectively support that this scorpion is a separately evolving lineage and should be recognized as a full species.

### 4.2 Unresolved phylogeny of *Mesobuthus caucasicus* complex

Although it has been demonstrated in the present study and in other studies that mtCOI DNA sequences are robust in defining species boundaries (Parmakelis et al. 2006; Shi et al. 2013; Fet et al. 2018), their power for resolving phylogenetic relationships among *Mesobuthus* scorpions appears limited. Given the several internodes of the phylogenetic tree are not well supported, the interspecies relationships among members of the *M. caucasicus* complex remain to be fully resolved in the future. Nevertheless, the clustering of the Chinese scorpion, *M. martensii*, within the clade of the species complex is strongly supported by both ML and Bayesian analyses (85/1.00). In both ML and Bayesian trees, a subclade of *M. martensii* clustered with subclades representative of the *M. caucasicus* complex in a major clade that was reciprocally monophyletic with respective to the mottled scorpion *M. eupeus* (Fig. 9). The mtCOI DNA sequences distances between *M. martensii* and *M. caucasicus* s. l. range from 7.2% to 8.4% which is significantly smaller than the genetic distances between some members (i.e. subspecies recognized earlier) of the *M. caucasicus* species complex (Table 2). As mentioned earlier, there is not clear geography barrier exist between *M. martensii* and *M. przewalskii*. We call the major clade includes the *M. caucasicus* complex and *M. martensii* collectively the *M. caucasicus* species group. Thus the *M. caucasicus* species group forms a phylogenetically and geographically coherent lineage that include 11 species spreading from the Caucasian region by *M. caucasicus* in the west, across the Tianshan Mountains, and to the East China Plain by *M. martensii*.

*Mesobuthus caucasicus* complex is particularly controversial in taxonomy. The earlier species of *M. caucasicus* has been placed in a separated monotypic genus *Olivierus* (Farzanpy, 1987) without solid justification or revision (Fet et al. 2000). However, this opinion was disputed by the first DNA phylogeny and the genus *Olivierus* was synonymized with *Mesobuthus* (Gantenbein et al. 2003). Recently, Kovařík (2019) split the genus *Mesobuthus* into three separate genera and resurrected *Olivierus* by putting all members of the *M. caucasicus* complex including the Przewalski’s scorpion, *M. martensii* and *M. mischi* as well as several other species (18 species in total) in it. We found Kovařík’s (2019) reassessment incurred even more controversial from a phylogenetic point of view. A quick contrast of Kovařík’s (2019) arrangement to recent phylogeny inferred from combined mitochondrial COI sequences and 16S sequences revealed that both the revised *Mesobuthus* and the resurrected *Olivierus* were not monophyletic (Fet et al. 2018). For example, *M. mischi* and members of *M. caucasicus* were included in *Olivierus* but they appeared polyphyletic in the mtDNA phylogeny with respective to *M. gibbous* and *M. cyprius*, both of which were assigned to a new genus *Aegaeobuthus* by Kovařík’s (2019). We acknowledge that the mitochondrial phylogeny also subjects to improvement and further validation. The highly similar genetic distances among species (Table 2) and poorly resolved phylogeny (Fig. 9-10) suggest that the *M. caucasicus* species complex might have undergone radiative speciation that gave rise to multiple species in a short time interval in early evolutionary history, possibly in response to the regional tectonic evolution and climatic changes (Shi et al. 2013). The quick succession of the speciation events means that the stochastic nature of the coalescent process in the ancestral species will cause different genes of the genomes to have difference genealogical histories, i.e. gene trees, due to incomplete lineage sorting of the ancestral polymorphisms (Shi & Yang 2018). Thus, it can be expected that gene trees inferred from other loci will not necessarily be identical to the existing mtDNA-based phylogenetic tree. Thus, it is prudent to keep the genus concept of *Mesobuthus* unchanged before a thorough and robust revision can be performed based on extensive sampling of taxa and characters (relevant to morphology, genetics and ecology) that reflect contingent properties of each species.

## Acknowledgements

We would like to thank Dr. Xianguang Guo for providing scorpion specimens and Dr. Ming Bai for use of photographic equipment. We are grateful to Dr. Zhiyong Di and two anonymous reviewers for their constructive comments and suggestions.

## Figures

**Figure S1.**
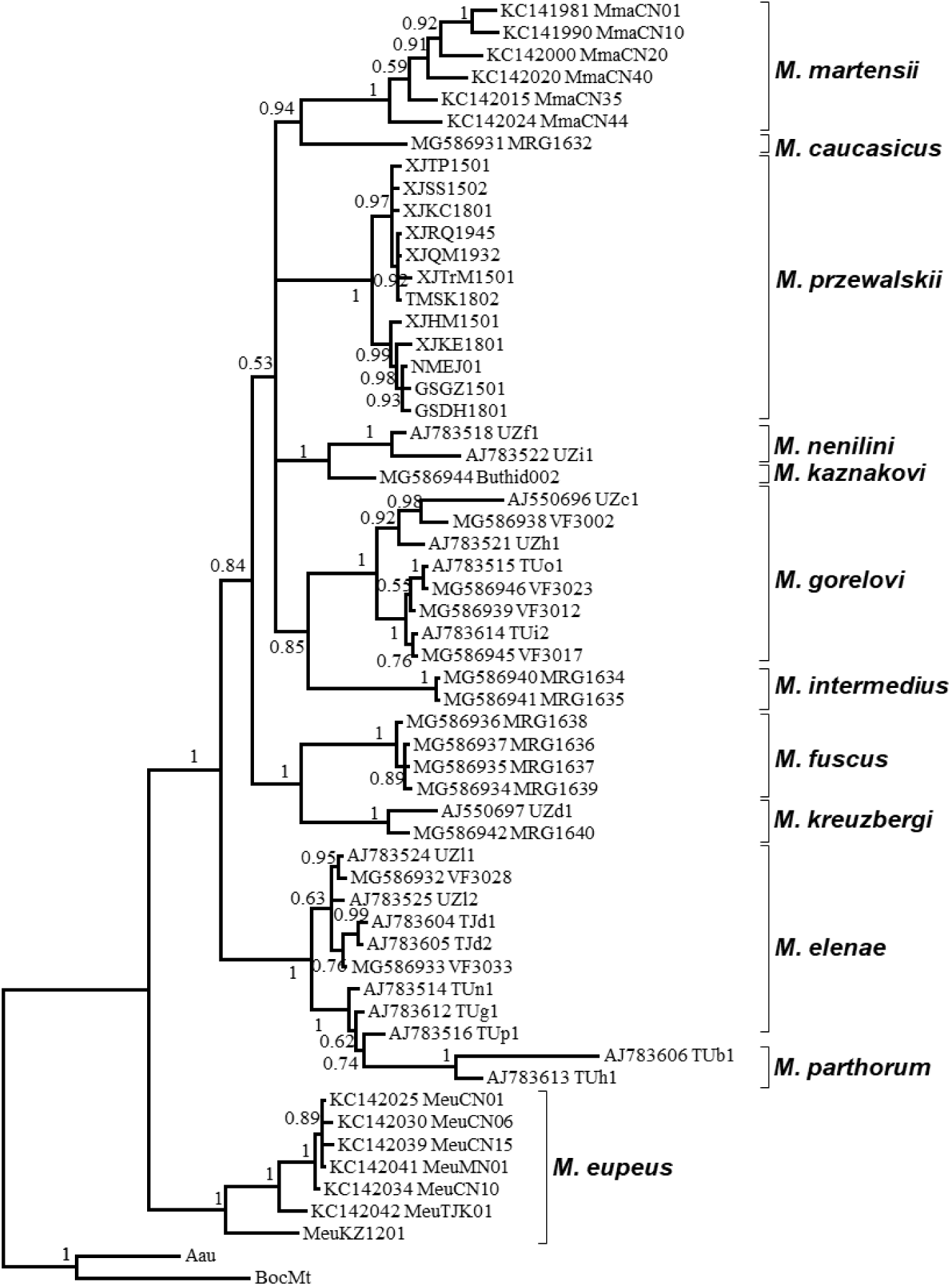
Bayesian consensus tree of the *Mesobuthus caucasicus* complex reconstructed using mitochondrial DNA sequences.

